# Refinement of Cryo-EM 3D Maps with Self-Supervised Denoising Model: crefDenoiser

**DOI:** 10.1101/2023.12.15.571622

**Authors:** Ishaant Agarwal, Joanna Kaczmar-Michalska, Simon F. Nørrelykke, Andrzej J. Rzepiela

## Abstract

Cryogenic electron microscopy (cryo-EM) is a pivotal technique for imaging macromolecular structures. Despite extensive processing of large image sets collected in a cryo-EM experiment to amplify the signal-to-noise ratio, the reconstructed 3D protein density maps are often limited in quality due to residual noise, which in turn affects the accuracy of the macromolecular representation. In this paper, we introduce crefDenoiser, a denoising neural network model designed to enhance the signal in 3D cryo-EM maps produced with standard processing pipelines, beyond the current state of the art. crefDenoiser is trained without the need for ‘clean’, ground-truth target maps. Instead, we employ a custom dataset composed of real noisy protein half-maps sourced from the Electron Microscopy Data Bank repository. Strong model performance is achieved by optimizing for the theoretical noise-free map during self-supervised training. We demonstrate that our model successfully amplifies the signal across a wide variety of protein maps, outperforming a classical map denoiser and a network-based sharpening model. Without biasing the map, the proposed denoising method often leads to improved visibility of protein structural features, including protein domains, secondary structure elements, and amino-acid side chains.

## 1 Introduction

### 1.1. Noise sources and denoising in cryo-EM

Cryogenic electron microscopy (cryo-EM) is one of the leading methods to elucidate protein structures (Bai et al., 2015; Cheng, 2018). In a cryo-EM experiment, a low-intensity electron beam must be used in order to minimize the organic sample degradation during imaging, resulting in noisy images. Thousands of these noisy images are collected and processed to improve the low, significantly below 1 (Frank and Al-Ali, 1975; Egelman, 2016), signal-to-noise ratio (SNR). This approach ultimately allows for modelling detailed, angstrom-resolution 3D protein maps. Still, the remaining noise is one of the factors limiting the reconstructed map’s quality (Rosenthal and Henderson, 2003; Frangakis, 2021).

The low-intensity electron beam is responsible for the shot noise on the cryo-EM images. Further, the protein particle projections are modulated by structural noise. This noise appears due to the non-uniform surroundings of the imaged particles, for example, amorphous ice impurities and ice thickness fluctuations. The resulting 3D protein density maps are also affected by errors in data processing, for instance, inaccuracies of 3D image alignment (Jiménez-Moreno et al., 2021).

The most successful cryo-EM denoising method so far is simply processing and averaging a large number of images. The improvement of the reconstructed map as a function of the number of acquired images can be estimated ((Rosenthal and Henderson, 2003)). The relation is logarithmic, which sets improvement limits due to acquisition costs. Other methods, such as 2D micrographs denoising (Palovcak et al., 2020; Bepler et al., 2020), 3D map denoising (Ramlaul et al., 2019; Tegunov et al., 2021), and corrections for 3D image alignment (Jimeénez-Moreno et al., 2021) are an active area of development and testing.

### 1.2. Deep learning enhances cryo-EM data processing

Deep learning has found extensive use in image processing. Therefore, the adoption of neural network models in cryo-EM processing is broad and the transfer of new methods has often been straightforward (Chung et al., 2022). For example, the popular YOLO object detection network (Redmon et al., 2016) was adapted to pick protein particles from EM images as crYOLO (Wagner et al., 2019). Denoising network models, developed for general-purpose image denoising and restoration, can also be used for contrast enhancement in the 2D EM images (Bepler et al., 2020; Lehtinen et al., 2018; Batson and Royer, 2019). There are also a number of specialized methods for 3D model building (e.g. CryoDRGN (Zhong et al., 2021), 3DFlex (Punjani and Fleet, 2023)) map postprocessing (DeepEMhancer (Sanchez-Garcia et al., 2021), EMReady (He et al., 2023)), map analysis (DeepRes (Ramírez-Aportela et al., 2019)) and atomistic model building (Emap2sec (Maddhuri Venkata Subramaniya et al., 2019), ModelAngelo (Jamali et al., 2022)), which are powered by neural networks.

### 1.3. 3D map sharpening and denoising

Cryo-EM 3D density maps show loss of contrast at high resolutions. This is caused by the decay of high-frequency signal amplitudes, which are smaller than expected when compared to the reference X-ray scattering data (Rosenthal and Henderson, 2003). Contrast degradation is caused by imperfect imaging due to limitations of TEM apparatus, including specimen movement and charging, radiation damage, inelastic electron-scattering events, partial microscope coherence, particle flexibility and heterogeneity, and also limitations of data processing methods (Henderson, 1992; Rosenthal and Henderson, 2003). To restore the degraded signal, global sharpening methods and, more recently, local sharpening methods have been developed. LocScale (Jakobi et al., 2017) uses an atomic reference structure to locally correct signal amplitudes. LocalDeblur (Ramírez-Aportela et al., 2020) performs deblurring based on the local resolution estimation. DeepEMhancer (Sanchez-Garcia et al., 2021) is a network model trained on pairs of raw experimental maps and post-processed, sharpened maps, which performs sharpening and denoising in one step. EMReady is a similar method (He et al., 2023) but trained on pairs of raw experimental maps and maps simulated from atomistic models. These two methods are optimized to match the ground truth final post-processed maps but not necessarily to represent the raw experimental data optimally. Another relevant method, LAFTER (Ramlaul et al., 2019) is a classical local 3D map denoising algorithm based on two serial filters. LAFTER operates in both real and Fourier space. It compares independent half-set reconstructions to identify and retain shared features with power greater than the noise. LAFTER does not sharpen EM maps but only denoises them, which makes it a suitable reference method for benchmarking our network-based denoising model.

### 1.4. Contributions

Entries in the Electron Microscopy Data Bank repository (EMDB) contain not only final processed cryo-EM 3D maps but, in many cases, ‘half-maps’, which result from processing two randomly divided half data sets. The two half-maps are used for the analysis of the map accuracy (Van Heel and Schatz, 2005; Rosenthal and Henderson, 2003) and can be used to illustrate how the map’s SNR changes in the function of the signal frequency with Fourier Shell Correlation plots (FSC, (Van Heel, 1987)) or by directly calculating power spectra of signal and noise components ((Palovcak et al., 2020)). This type of data is suitable for training a neural network 3D map denoising model. The most natural setup would be the so-called noise-to-noise model (Lehtinen et al., 2018), in which the first half-map is used as a denoising template, and the second serves to calculate loss during the supervised model training. This is how the M software (Tegunov et al., 2021) is, on the fly, training a map-specific model during a map refinement procedure. Here, we take advantage of existing theoretical analysis to further enhance the model’s denoising power. Rosenthal et-al. derive a relation between an ideal, noise-free 3D map and a pair of two noisy half-maps as a function of FSC (Rosenthal and Henderson, 2003). In the Methods Section 2, we outline how we employ this relation to optimize the denoising network in self-supervised training. We compare our model crefDenoiser to the recent 3D map denoiser, LAFTER (Ramlaul et al., 2019) and sharpening model EMReady (He et al., 2023), and we show better denoising performance on the analyzed maps sets. Further, for the test set, we analyze the signal-to-noise enhancements in the function of signal frequency, and we show that crefDenoiser does not introduce any large biases to the denoised maps. We also compare the denoised low-resolution maps to the higher-resolution maps to show denoising improvements. Finally, we provide examples of denoising with selected maps, where our processing provides insights into the usability of denoising.

## 2. Methods

### 2.1. Model Optimization

crefDenoiser is trained using a loss function based on the Fourier Shell Correlation and the statistical measure known as C_ref_. FSC score is the most popular metric used in cryoEM imaging to determine image and map quality (Rosenthal and Henderson, 2003). It measures the normalized cross-correlation between two volumes over corresponding shells in the Fourier domain. It quantifies the similarity of signals between two maps (or images when a 2D signal is analyzed) as a function of frequency. The FSC value between two map volumes is given by:

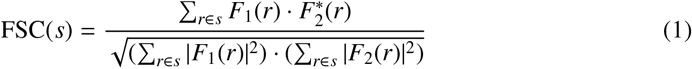

where *F*_1_ and 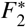 represent the Fourier transform and conjugate Fourier transform of the two volumes and *s* is the shell being considered. The summation is performed over all frequency voxels *r* contained in the shell *s*. To calculate a scalar score, we integrate the FSC curve over all frequency shells up to the Nyquist frequency.

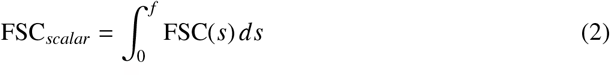

FSC values can range between +1 for perfectly correlated images to 0 for completely uncorrelated images. Negative values (up to −1), imply a negative correlation. An FSC of −1 would represent identical images with opposite contrasts (Penczek, 2020). In cryo-EM imaging, the FSC_half_ curve between half-data set maps is used to determine the so-called “gold-standard” resolution (Rosenthal and Henderson, 2003). The frequency at which the FSC curve first falls below a fixed value (usually 0.143) (Rosenthal and Henderson, 2003) is used as a resolution estimate.

*F*_1_ and *F*_2_ in Equation 1 can be represented by a common signal term and an additional noise term, *F*_1_ = *S* + *N*_1_, *F*_2_ = *S* + *N*_2_, where *N*_1_, *N*_2_ are realizations of noise *N*. With this, FSC_half_ becomes (Rosenthal and Henderson, 2003)

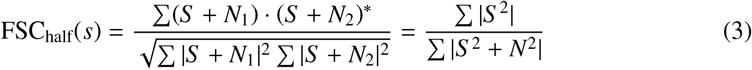

when signal and noise are uncorrelated and data in the half-sets is on the same scale. Using the above notation, we can also write FSC between an ideal map and a map reconstructed from a complete data set. The ideal map has no noise term, and the noise of the full-dataset map becomes 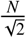, when compared to the half-data set noise, *N*. This so-called C_ref_ can be expressed in function of FSC_half_ (substituting with the result of Equation 3, see also (Rosenthal and Henderson, 2003)):

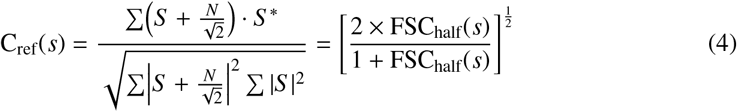

Our loss function is the mean absolute error between the C_ref_ in Equation 4 (calculated using readily available FSC_half_ curves), and FSC_FD_, which is calculated between the average of halfmaps (used as the network input map for denoising) and the denoised output map.

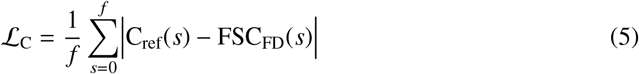

Calculating the loss, we assume that the average of two half-maps represents a map reconstructed from a complete data set.

The FSC_FD_ between a noise-free map and the average of two half-set maps should completely overlap with C_ref_. An FSC_FD_ value above C_ref_ indicates that the denoised map still contains some residual noise (under-denoised), while a value below C_ref_ points to a loss of signal (over-denoised). The lower the ℒ _C_, the closer the FSC_FD_ of our network output is to the C_ref_, indicating a more effective denoising operation. ℒ _C_ is differentiable since it is directly derived from the FSC function, which is differentiable (Kaczmar-Michalska et al., 2022) and can be readily applied in a gradient-based model training. The ℒ _C_ loss allows to perform Fourier-space-based model optimization for the real space, theoretical noise-free map, even without actually having noisefree maps to drive the model training.

### 2.2. Bias Analysis

The denoising process might introduce a spurious bias signal to the map (Palovcak et al., 2020). A noisy map consists of signal and noise:

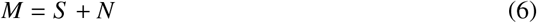

and a denoised map consists of signal, bias, and some leftover noise:

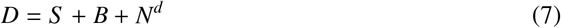

A variance of signal var(*S*), noise var(*N*), bias var(*B*), and leftover noise after denoising var(*N*^*d*^), can be calculated from the noisy and denoised half maps. We follow derivations provided in Palovcak et al. (2020). Elementary relations between variance and covariance of variables are used in the calculations and provided in the Supplementary Methods section 7.1. Assuming that *S* and *N* are independent, variance of signal and noise in 3D cryo-EM maps, var(*S*) and var(*N*), can be calculated using the noisy half-maps *M*_1_, *M*_2_:

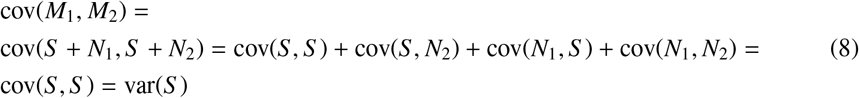

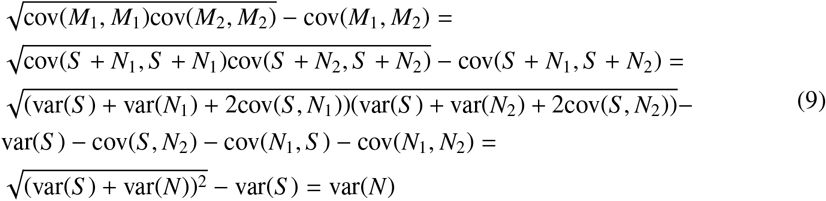

Further, assuming that *B* and *N, B* and *N*^*d*^, and *S* and *N*^*d*^, are independent, we can calculate the variance of *B* using the noisy, *M*_1_, *M*_2_, and denoised, *D*_1_, *D*_2_, half-maps.

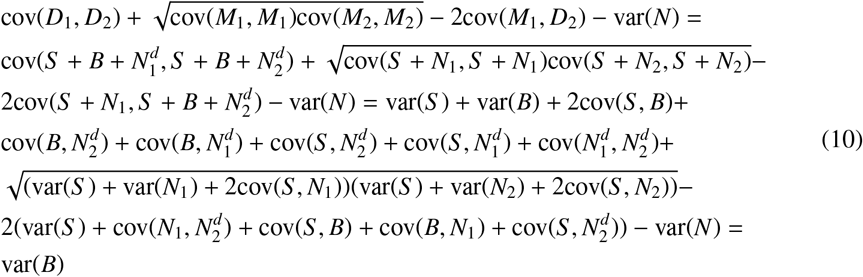

A similar derivation for *N*^*d*^ is provided in the Supplementary Methods. The covariances can be calculated separately for each frequency shell *s* in 3D maps:

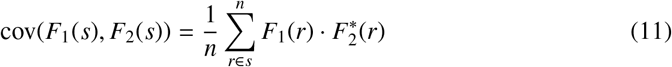

where *n* is the number of voxels *r* in the shell *s*.

### 2.3. Network Architecture

We use a 3D UNet-like (Ronneberger et al., 2015) model with five levels of depth in the contracting path and corresponding five levels in the expanding path. The overall architecture of crefDenoiser is enumerated below:

1. Input Layer: The model takes as input a 3D image with a single channel.
2. Contracting Path: The contracting path consists of five blocks, each containing a 3D convolutional layer with 16 filters, a Leaky ReLU activation function, and a 3D MaxPooling layer.
3. Bottleneck: The bottleneck consists of a 3D convolutional layer and a Leaky ReLU activation function. This part of the network is responsible for learning the most abstract features of the input data.
4. Expanding Path: The expanding path also consists of five blocks, each containing a 3D UpSampling layer, a concatenation operation, two 3D convolutional layers, and two Leaky ReLU activation functions. The concatenation operation combines the features learned in the contracting path with the upsampled output, allowing the network to use both local and global features for the reconstruction of the denoised image.
5. Output Layer: The final layer of the network is a 3D convolutional layer with a single filter, which outputs the denoised 3D image.

The total number of parameters in the model is 322,881, all of which are trainable.

Since cryo-EM images inherently capture the intricate three-dimensional structures of macromolecules, this architecture is particularly well suited to the task due to its ability to effectively learn spatial hierarchies and extract features from the 3D data. Although the network is trained on patches of maps (refer section 2.4), it is fully convolutional and can denoise whole maps of any size without any architectural restrictions.

### 2.4. Data Preparation

Our model is trained on data collected from the EMDB repository. All cryo-EM entries, with an associated mask and two half-maps attached, were downloaded from the online EMDB FTP server. Any entries with size mismatches between the two half-maps and/or the mask files were pruned. The remaining 3710 records, with resolutions in the range (1.22Å9.9Å), were used in constructing the train and test datasets.

All the half-map pairs were first independently masked and standardized to have a mean voxel value of 0 and an intensity standard deviation of 1. They were then randomly shuffled (as a pair) and split into patches of size 96×96×96. Any patches which lay completely outside the masking region were removed. Those remaining were then divided in a 1:9 ratio to construct test and train datasets. The training set finally contains 55176 such pairs of half-map patches from 3386 maps, while the test dataset contains 6126 pairs from 324 maps.

### 2.5. Training Process

The training process was conducted using the ADAM(Kingma and Ba, 2014) optimizer (*β*_1_ = 0.9, *β*_2_ = 0.999, *ϵ* = 10^−8^) with an initial learning rate of 0.0003. The learning rate was reduced exponentially with a decay rate of *k* = 0.7 every ten epochs. The model was trained for 195 epochs with a batch size of 6 on three NVIDIA A100 80GB GPUs. The training time was approximately 120 hours. After each epoch, the model’s performance was evaluated on the validation set, and the model weights were saved.

### 2.6. Map Sharpening

cREF-denoised maps may be further sharpened. Here the selected denoised maps were sharpened to facilitate graphical comparison with the published maps. Local sharpening with Phenix software (Adams et al., 2010), with the resolution threshold set to be slightly lower than the published resolution (−0.5 Å) was chosen in the Phenix setup. The sharpening method was automatically selected by Phenix.

## 3 Results

### 3.1. Comparison to LAFTER and EMReady

For the set of masked denoised test maps, we have calculated FSC_FD_ and compared it to the theoretical C_ref_. Figure 1 **A**, shows RMSE results for denoising with crefDenoiser and the LAFTER denoiser. The test set averaging is performed in frequency bins up to a map resolution (C_ref_ = 0.5), meaning that the RMSE values in the high-resolution bins are calculated with fewer data points (i.e., maps) than in the low-resolution bins since not all the tested maps are high resolution. We observe that crefDenoiser-generated models have a particularly low deviation from the ideal model for low-resolution signal, and for resolutions higher than 10 Å, the deviation is roughly constant but about 1-2 orders of magnitude higher. The frequency averaged RMSE equals 0.05. With C_ref_ always higher than 0.5, the mean error is on average less than 10 percent of the correlation signal. The performance of the LAFTER denoiser is significantly worse, with frequency averaged RMSE equal to 0.78. The distribution of LAFTER average RMSE results is bimodal, as seen in Figure 1 **B**, with peaks around 0.25 and 0.8. We observe that LAFTER can effectively denoise only a fraction of the tested maps. The distribution of frequency-averaged RMSE results for crefDenoiser is unimodal, with the highest average RMSE values around 0.25 (Figure 1 **B**). In Figure 2 the FSC_FD_ and C_ref_ for one map, EMD 11698, is shown. FSC_FD_ follows closely the theoretical C_ref_ curve. This is the case also for the other tested maps.

**Figure 1:**
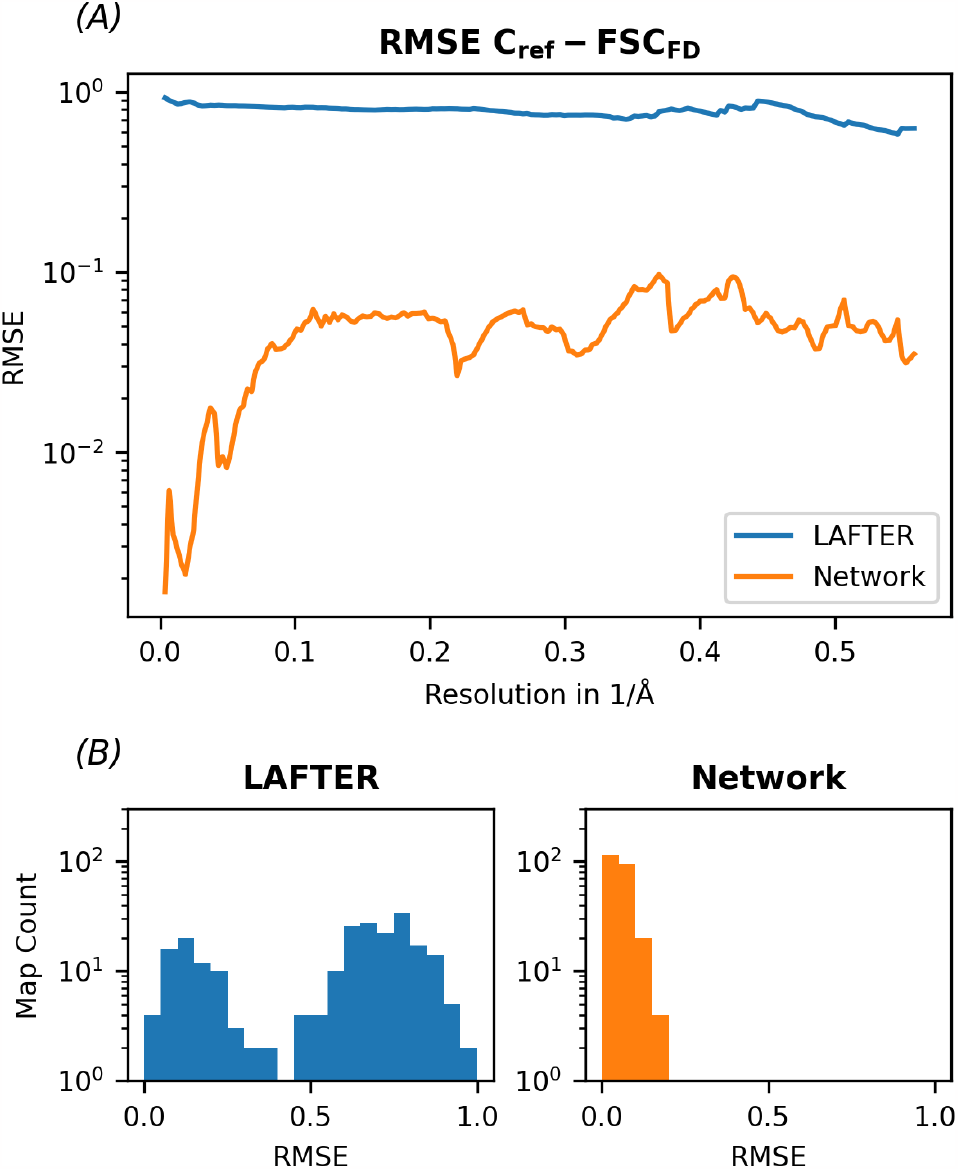
Analysis of denoising performance for the test set of 3D cryo-EM maps. Root mean square error, RMSE between FSC_FD_ and C_ref_ is plotted for frequency binned data in panel **A** for the network denoised and LAFTER denoised maps. The averaging is performed in frequency bins up to a map resolution (C_ref_ = 0.5), meaning that the RMSE values in the high-resolution bins are calculated with fewer data points than in the low-resolution bins. In panel **B**, histograms of frequency-averaged RMSE values are shown.

**Figure 2:**
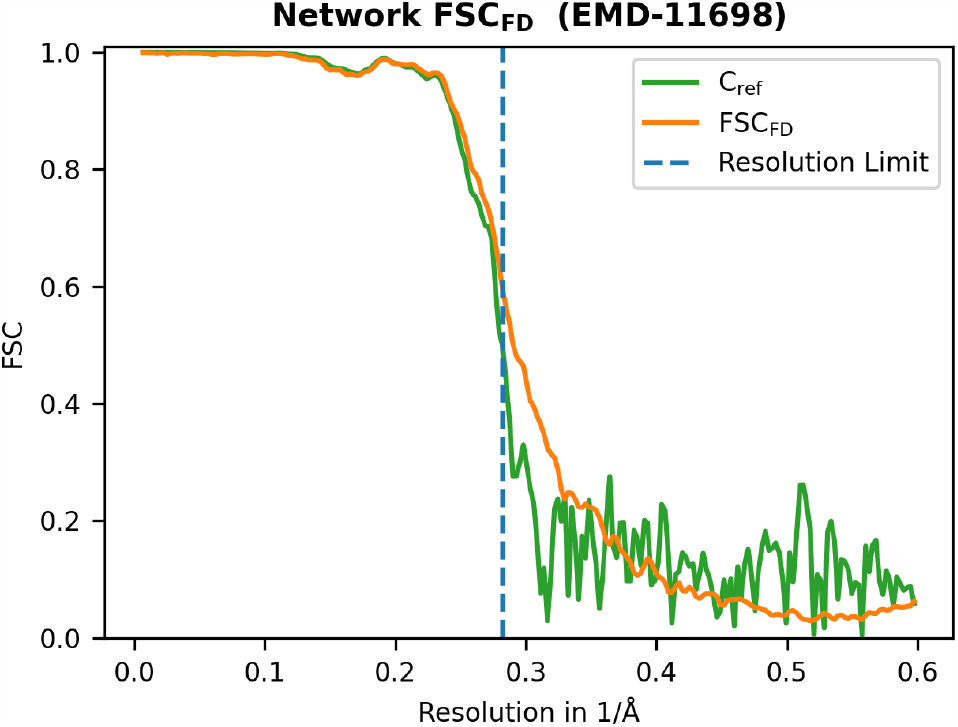
Analysis of denoising performance for a single cryo-EM map, PspA helical assembly (EMD 11698). FSC_FD_ and C_ref_ are plotted up to the Nyquist resolution. The dotted line marks the map resolution.

Further, we compared denoised maps to maps processed with the EMReady (He et al., 2023) model. EMReady authors use the resolution at which the FSC between a pair of maps falls to one-half (i.e., FSC-0.5) as a metric for the model evaluation. We compare our performance using this same metric. For this, we applied crefDenoiser on a set of 25 half-map pairs that were used in the analysis presented in Figure S2 of the EMReady manuscript. We note that these maps belong to both our model’s training and test set. Next, we calculated FSC−0.5 values for the noisy (FSC−0.5(M_1_, M_2_)), denoised-noisy half-map pairs (FSC−0.5(D_1_, M_2_), FSC−0.5(D_2_, M_1_)) and EMReady processed maps 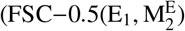, 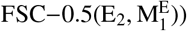. In Figure 3 and Figure S1, we present the change of FSC − 0.5 after applying crefDenoiser and the EMReady model. We note that FSC − 0.5 values, published in the EMReady supplementary dataset, between noisy half maps 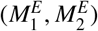, are slightly different when compared to the values available in the EMDB repository (or calculated by the EMDA (Warshamanage et al., 2022) package with the noisy maps, FSC − 0.5(M_1_, M_2_)), especially for lower resolution maps. This is likely due to the applied preprocessing in *M*^*E*^ maps, including voxel size normalization. For crefDenoiser we do not normalize voxel size before map denoising, and therefore we use original noisy maps in the analysis. For most of the analyzed maps, the FSC − 0.5 change is negative for both methods. Such change suggests improved quality of the processed maps since the FSC − 0.5 shifts to higher resolution values. Even though EMReady was not trained to denoise 3D density maps, we see a similar effect as for our model; however, crefDenoiser performs, on average, better than EMReady (Figure 3), irrespective of the map resolution (Figure S1). The presented analysis suggests that our denoising model can accurately denoise half-maps, even though it was trained on the full-data density maps.

**Figure 3:**
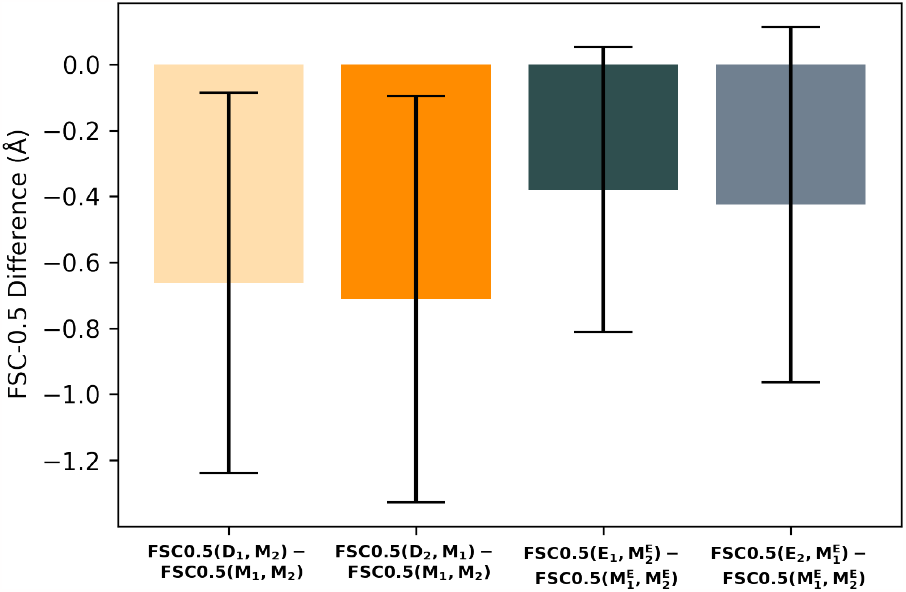
Comparison of FSC − 0.5 for maps denoised with our model (*D*_1_, *D*_2_) and EMReady network (*E*_1_, *E*_2_). The difference of FSC − 0.5 values for denoised-noisy half-maps to FSC − 0.5 of noisy-to-noisy half-maps, *M*_1_, *M*2, for 25 EMDB entries is shown. The analysis uses data presented in the Figure S2 of the EMReady manuscript, where FSC0.5 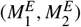 is slightly different than FSC-0.5(*M*_1_, *M*_2_), apparently due to preprocessing. Means and standard deviations for the set of analyzed maps are shown.

### 3.2. Comparison to published maps

Figure 4 compares lower-resolution denoised maps to high-resolution map. The analyzed apoferritin entries EMD-20026(1.8Å), EMD-20027(2.3Å), and EMD-20028(3.1Å) are reconstructed from the same data set using different fractions of the acquired single-particle images(Pintilie et al., 2020). Denoising improves FSC curves for both lower-resolution models. Mainly, the high-frequency part of the FSC curve is changed. While globally, the denoised maps more closely resemble the higher-resolution map we further analyzed if the denoising introduces any spurious densities. In Supplementary Figure S2, we show three areas of apoferritin and compare the deposited (EMD-20026,EMD-20027,EMD-20028) and denoised maps (EMD20027,EMD-20028) with atomistic model 3ajo. The network-denoised maps are sharpened with Phenix, to account for the missing contrast (see also: section 2.6). We have not detected model hallucinations in the denoised maps, while lower-density areas (side chains, ions) are slightly better represented.

**Figure 4:**
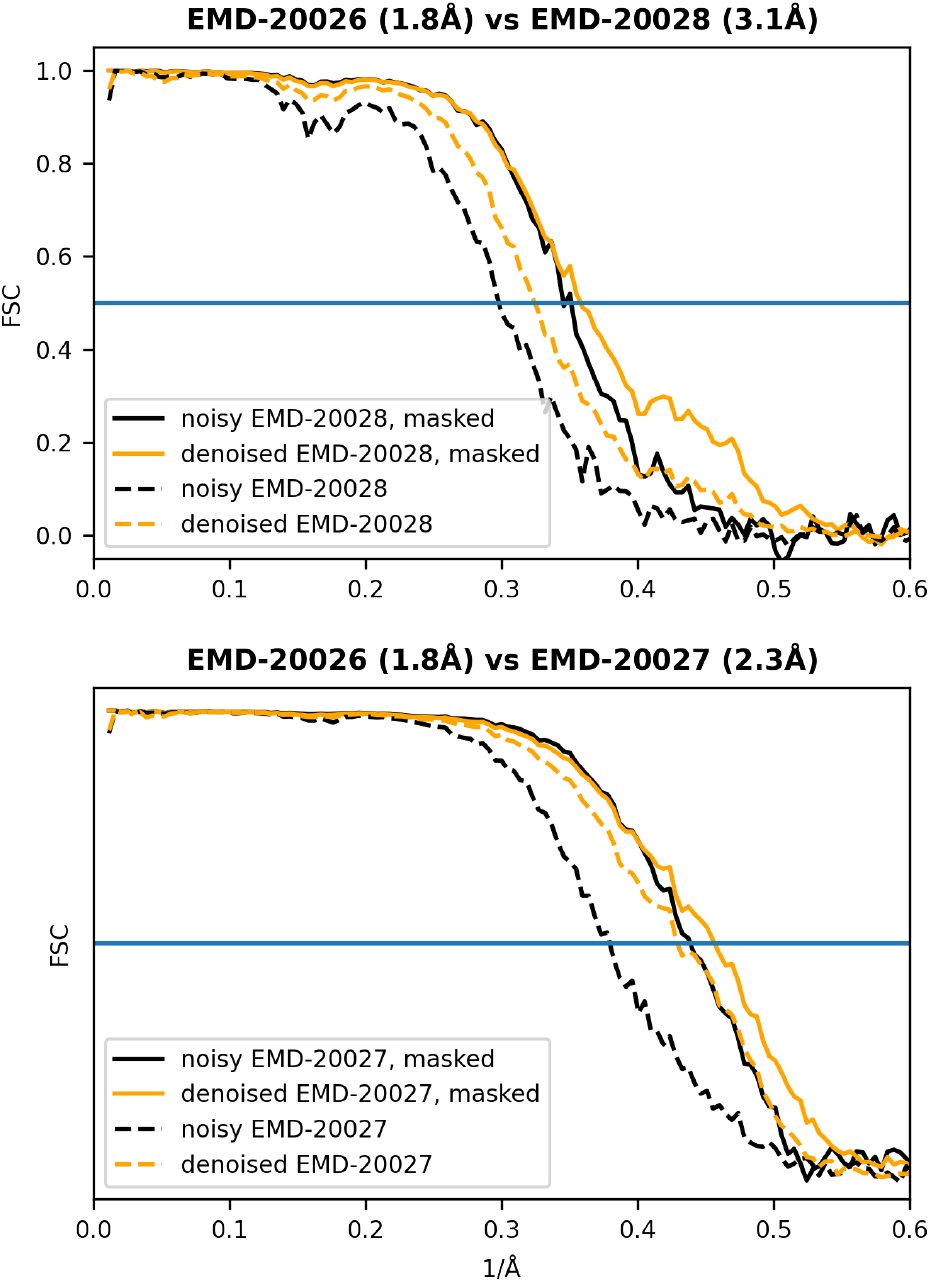
Comparison of denoised maps to a higher resolution map. Half-maps for human apoferritin EMDB entries EMD-20027 (2.3Å) and EMD-20028(3.1Å) were averaged (within entry), and the mean maps were subsequently denoised. Next, FSC curves to the mean map of the high-resolution entry, EMD-20026(1.8Å), were calculated. Maps EMD-20026, EMD-20027, and EMD-20028 were obtained by processing different fractions of the same single-particle data set(Pintilie et al., 2020). Denoising improves FSC curves to the higher resolution map for both masked and nonmasked maps. The maps are part of the test set.

In Figures 5 and 6 we visually compare crefDenoised maps to the published final 3D cryo-EM maps, as deposited by the authors in the EMDB repository. For example, in medium-resolution map EMD-22778(4Å) of Sec61 membrane channel(Itskanov et al., 2021) densities of some *α*helices improve after denoising (see Figure 5) while noise due to lipid densities is mostly removed. To exclude that the effects are only due to sharpening we also show noisy-sharpened maps. In the case of SARS-CoV-1 Spike protein map, EMD-34420, shown in Figure 6 denoising removes high-frequency noise that obscures overview of the domain positions within the spike map (Figure 6 D), while the high-resolution information, in particular densities of amino acid side chains, is unaffected (Figures 6 B and D).

**Figure 5:**
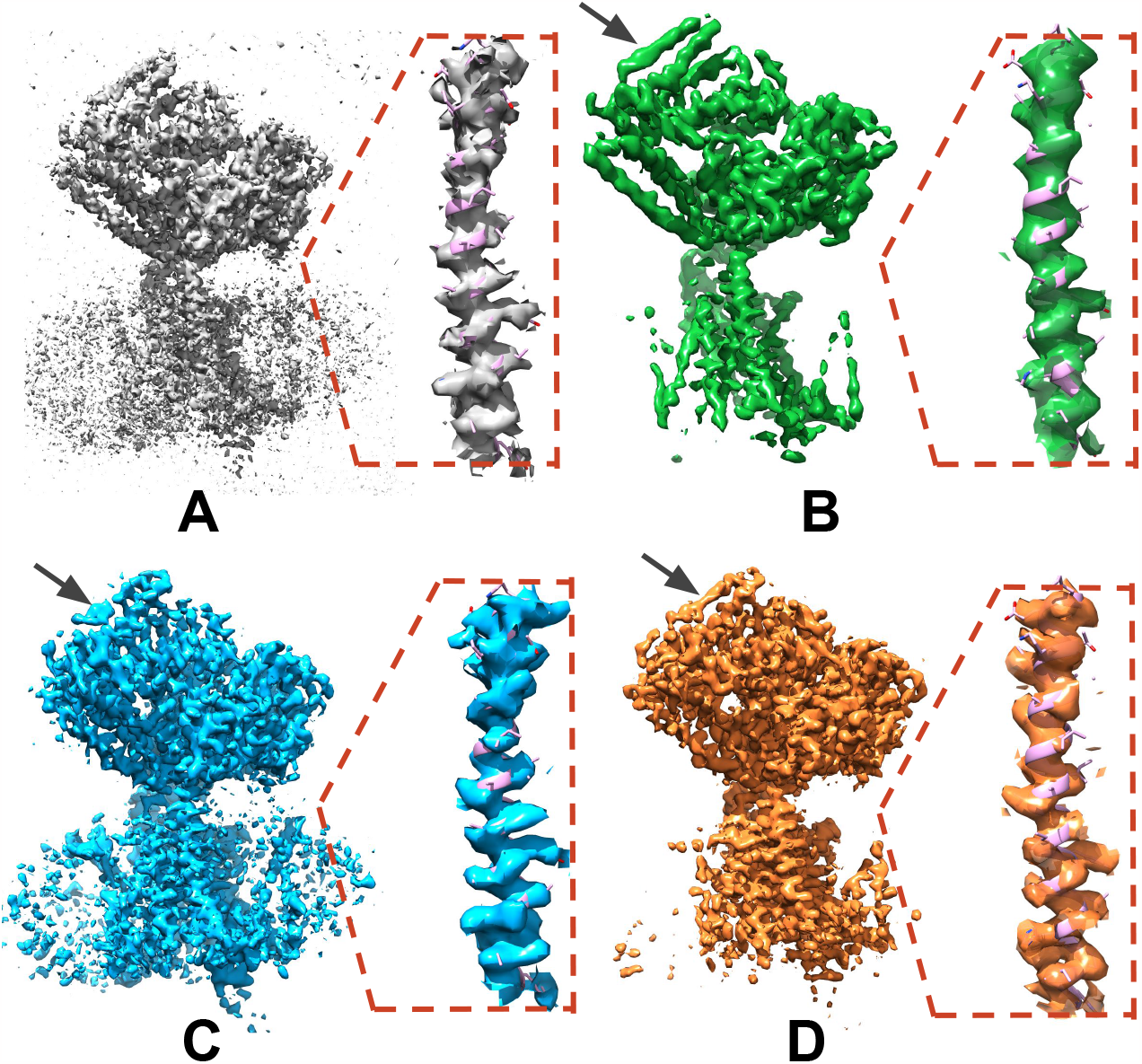
Denoising medium resolution map. The map of Sec61 channel from Saccharomyces cerevisiae (Itskanov et al., 2021), and its fragment focused on one selected internal channel helix with the fitted atomistic model are visualized. The map is part of the test set and has a resolution of 4 Å. The noisy map, constructed as a mean of two half maps, is presented in panel **A**. The final published map is shown in panel **B**. The noisy sharpened mean map is presented in panel **C**. The denoised and sharpened mean map is presented in panel **D**. The denoised map preserves lower-resolution motifs (black arrow marked *α*-helix) and high-resolution details (the insight) while the noise is substantially reduced. Contouring was tuned to make the maps most similar.

**Figure 6:**
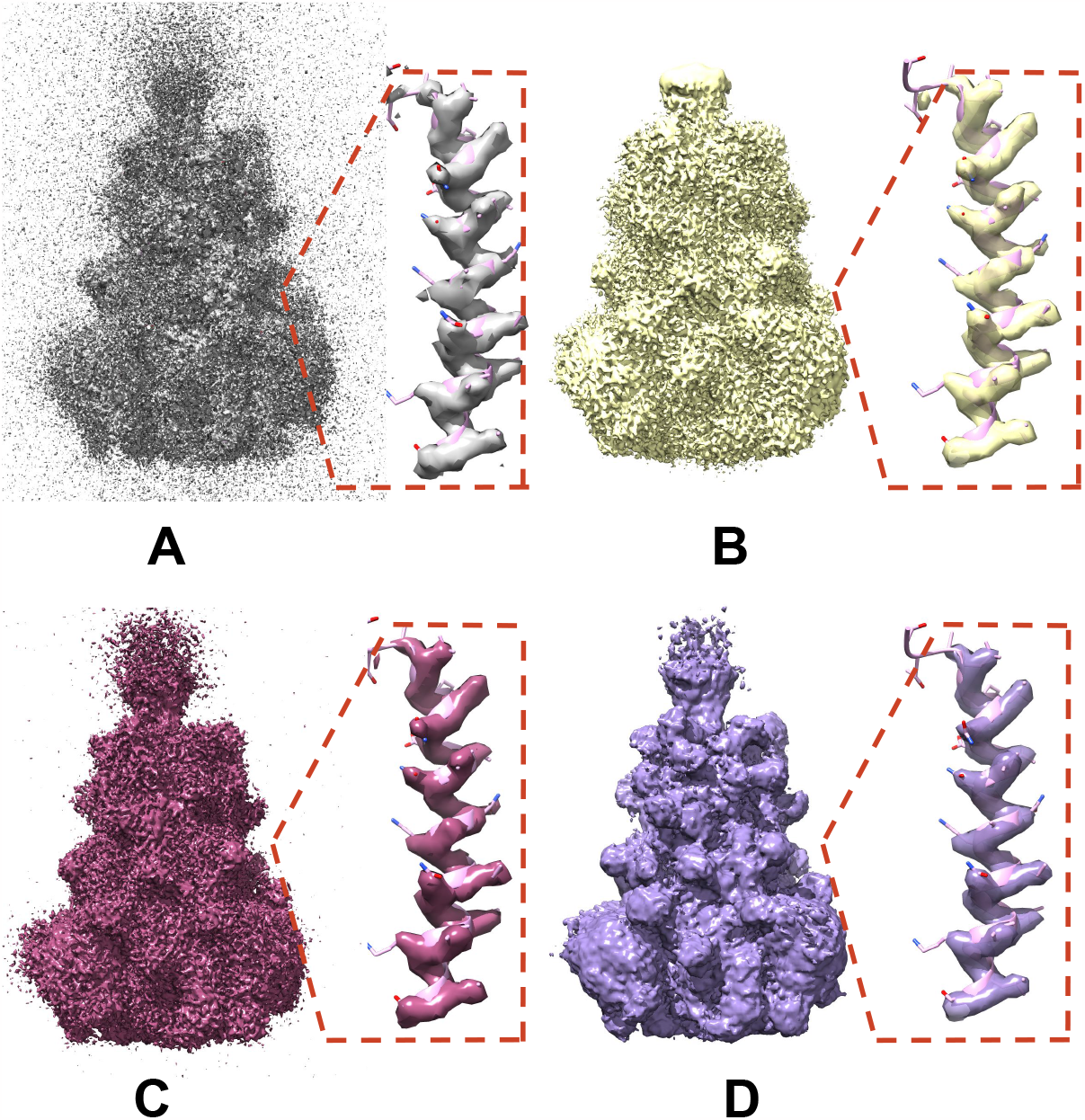
Denoising high-frequency noise. The map of SARS-CoV-1 Spike Protein (EMD-34420, to be published), and its fragment focused on a single helix with the fitted atomistic model are presented. The map is part of the train set and has a resolution of 2.99 Å. The noisy map, constructed as a mean of two half maps, is presented in panel **A**. The final map published in the EMDB repository is shown in panel **B**. The noisy sharpened mean map is presented in panel **C**. The denoised and sharpened mean map is presented in panel **D**. Contouring was tuned to make the maps most similar. The high-frequency structural features of the published and denoised maps are similar (see the *α*-helix), however, the denoised map provides a clear outlook of the overall architecture of the spike due to the removal of high-frequency noise.

### 3.3. Signal, noise and bias in the denoised maps

In Figure 7, we analyzed signal, noise, and bias components in the set of 100 denoised test half-maps. The separate components are computed as explained in section 2.2. The presented analysis assumes that crefDenoiser can also accurately denoise half-maps, even though it was trained on the full-data density maps. We reason that the noisy maps from the large train set have a broad range of noise levels, and most of the half-maps fall within that range. With this assumption, the denoising model should denoise a half-map like any other map with a similar noise level. In Figure 7, we show that, on average, the medium and high-frequency noise is significantly reduced in the denoised maps. While bias is prominent in the first several low frequencies, overall, it is much smaller than the signal. Only around 0.35 1*/*Å does the bias and leftover noise power spectrum cross with the signal power. In the noisy maps, noise and signal power spectrum cross around 0.27 1*/*Å, which suggests improved quality of the denoised maps.

**Figure 7:**
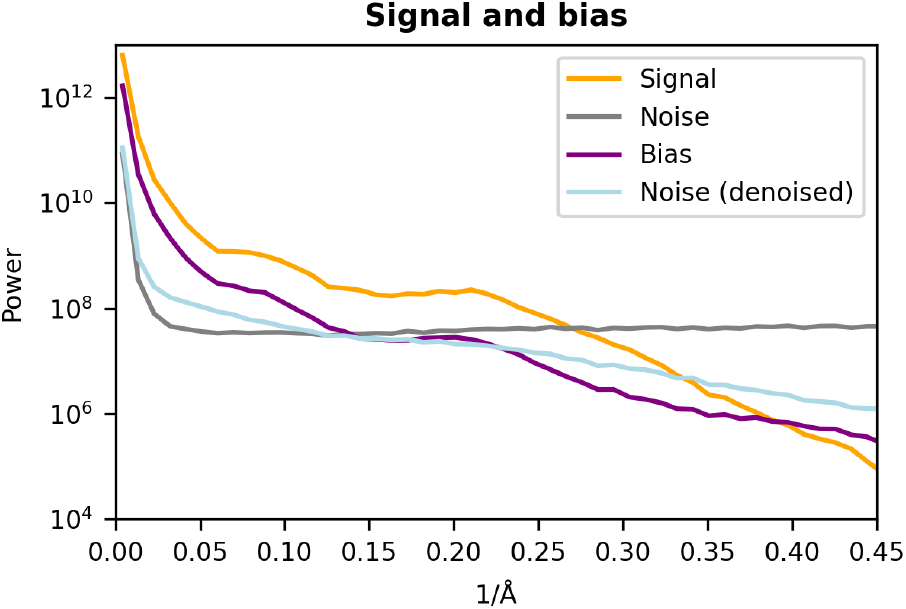
Denoising bias. The variance of signal (var(*S*)), noise (var(*N*)), bias (var(*B*)), and leftover noise after denoising (var(*N*^*d*^)) is plotted in the function of resolution. The mean values were calculated with 100 masked maps taken from the test set. Noisy half-maps and denoised half-maps were used to calculate the plotted characteristics as described in the text.

## 4 Discussion and Conclusions

This contribution presents crefDenoiser, a self-supervised deep network model for denoising 3D cryo-EM density maps. The network denoiser has better accuracy when compared to the recent classical denoiser, LAFTER, and to our knowledge, it is the first network-based model for denoising this type of 3D EM data in a self-supervised manner. The model is trained on about 3700 experimental maps deposited in the EMDB repository to optimize an ideal noise-free map using the presented theory-based loss. We showcase the benefits of denoising in 3D EM map analysis on several real-map examples, and we provide further data that confirm the improved quality of the denoised maps.

The recent sharpening models, DeepEMhancer (Sanchez-Garcia et al., 2021) and EMReady (He et al., 2023), might, as a side effect, also perform map denoising since they are trained to match post-processed and simulated maps in which noise levels are low. The presented crefDenoiser, on the other hand, is trained to maximize the consistency of raw cryo-EM data, i.e. half-maps, with the denoised map employing the C_ref_ based analytical loss. The output from crefDenoiser can still go through processing steps such as sharpening that are not particularly affected by denoising. As we indeed show, the EMReady model also improves FSC-0.5 values for the set of analyzed maps, but to a lower extent than the presented model.

Compared to a standard ℒ _2_ loss the C_ref_ based loss is more sensitive to high-frequency signal, which seems to be beneficial for model training. ‘Noise-to-noise’ models, where one 3D halfmap is used as denoising input, and the second half-map is used to compute loss, could also be trained with a high-frequency focused loss, such as FSC (equation 1, (Kaczmar-Michalska et al., 2022), see also (Tegunov et al., 2021)), instead of ℒ _2_ loss. However, the ‘noise-to-noise’ setup, as used for example in the M software (Tegunov et al., 2021) does not take advantage of the theoretical noise-free map. In our model, we use the mean of both half-maps as input for denoising training, with the noise magnitude reduced by 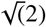. Further, the model is extensively trained on a relatively large data set (compared to M) until convergence, resulting in a versatile, high-performance denoiser.

While in most of the presented analysis, our results were calculated for maps that were not part of the training set, it is important to highlight that in real-world applications, the maps requiring denoising could also be incorporated into the training process. This flexibility is made possible by our model’s ability to train without the need for ground truth clean maps (self-supervised learning), and has the potential to yield even better denoising results, as the model can adapt to more specific noise patterns present in these maps.

In the ideal scenario, the denoised network generates an optimal denoised map. The current model, similar to models trained on a standard L_2_ loss (see (Menon et al., 2020)), generates a ‘mean’ solution, here a 3D density map, that approximates all possible noise-free maps. Such a ‘mean’ solution was shown to be of lower quality in the high-frequency domain due to bias (Palovcak et al., 2020). However, we observe that the leftover noise (see Figure 7), rather than the bias, is currently the main limiting factor of the denoised maps. Interestingly, the lower-resolution maps shown in Figure 4 show the highest improvement after denoising for frequencies where the collective change of signal-to-noise-and-bias ratio is the highest (Figure 2.2). Training models with a larger training data set, when it becomes available, might further reduce the leftover noise and improve the denoising results.

## 5. Code

The code and model are available for reviewers in the Github repository.

## 6. Acknowledgements

This work was performed with the support of the Swiss National Science Foundation SPARK grant nr CRSK-3-190804 to Andrzej Rzepiela and Novartis FreeNovation grant to Andrzej Rzepiela and Simon F. Nørrelykke. We would like to thank Daniel BÖhringer for discussing the manuscript.

## 7. Supplementary Methods

### 7.2. Variance and covariance relations

Having variables *X, Y, V*, and *Z*:

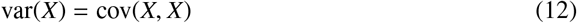

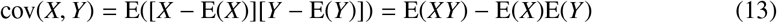

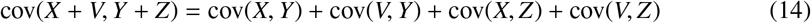

Equations 12 and 14 imply that

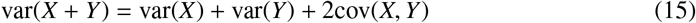

and if *X, Y* are independent, E(*XY*) = E(*X*)E(*Y*), equation 13 implies that cov(*X, Y*) = 0.

### 7.2. Derivation of var(N^d^)

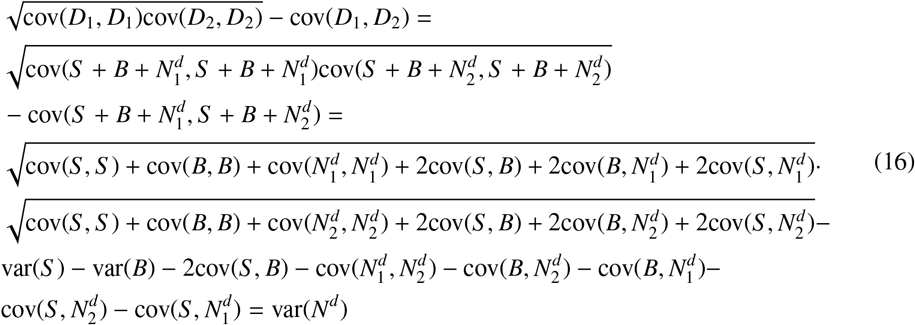

## 8. Supplementary Figures

**Figure S1:**
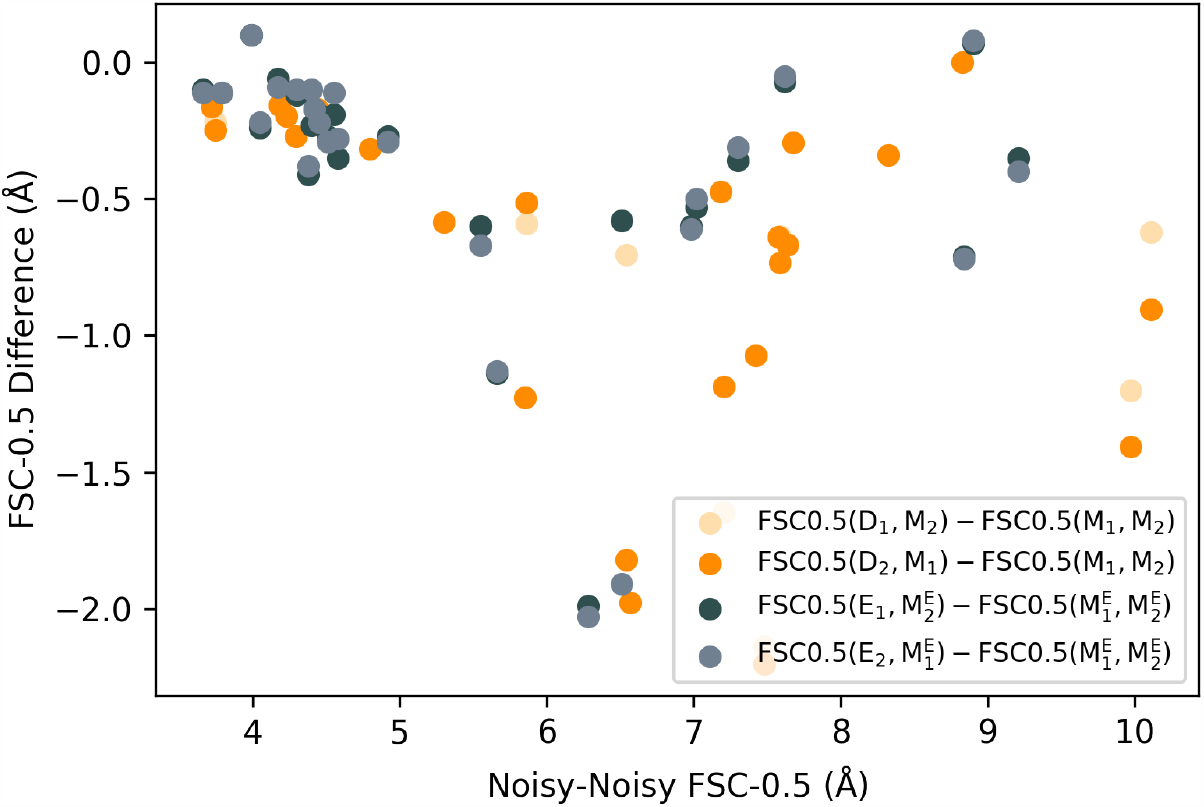
Comparison of FSC − −0.5 change after denoising for maps processed with our model (*D*_1_, *D*_2_) and EMReady network (*E*_1_, *E*_2_). The plot corresponds to data shown in Figure 3, but each map is shown as a separate point, without averaging over the whole set. The x-axis shows the FSC−0.5 for maps before network processing.

**Figure S2:**
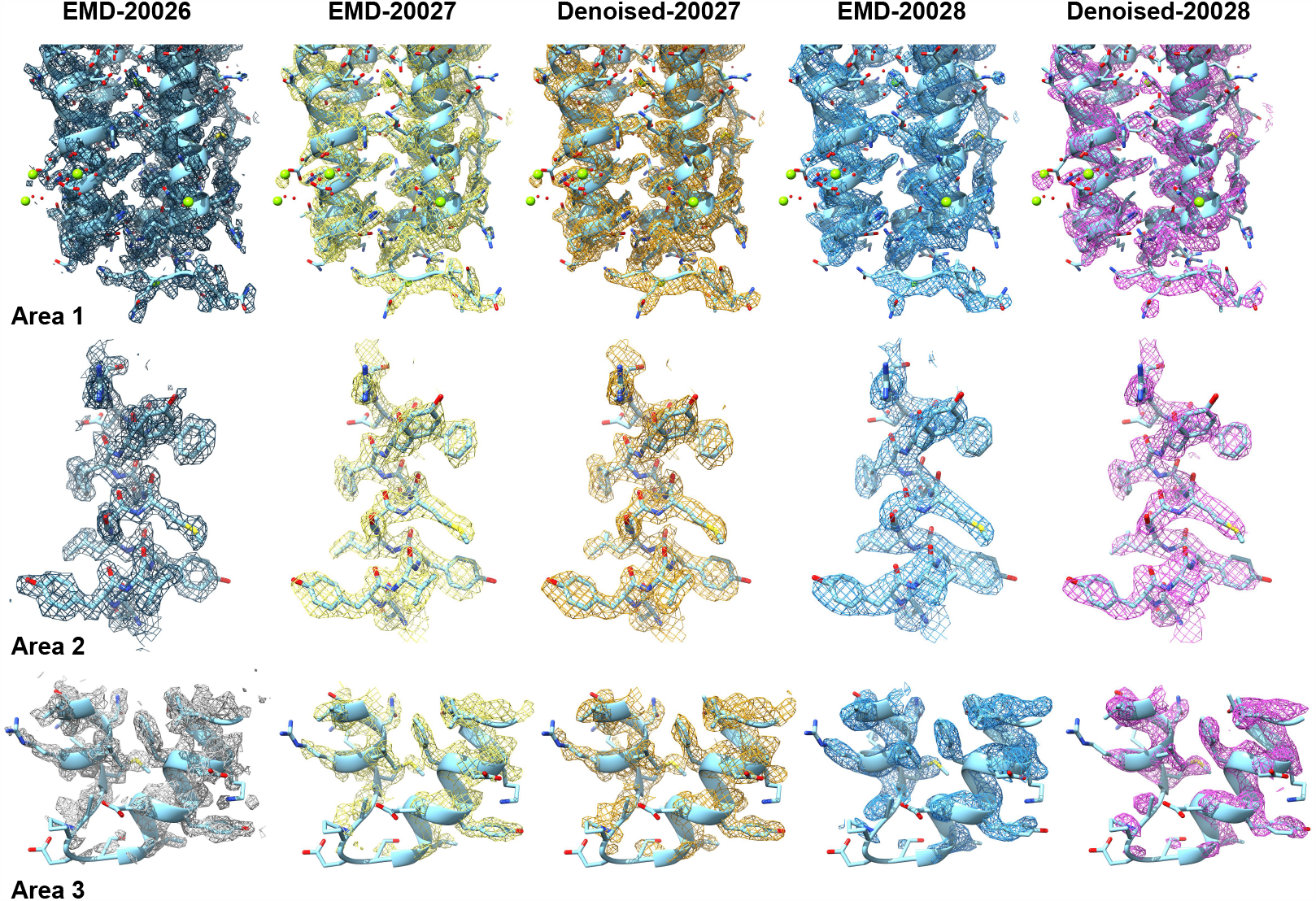
Denoising does not introduce noticeable false densities to the maps. Comparisons of three areas for the published human apoferritin maps EMD-20026(1.8Å) EMD-20027(2.3Å) and EMD-20028(3.1Å) and two denoised maps (from half maps of sets EMD-20027 and EMD-20028). The denoised maps were sharpened before comparison, as specified in section 2.2. Atomistict model (1.5Å) of apoferritin, 3ajo, is also shown. Contouring is the same for the three areas.

